# Mutation rate estimation for 15 autosomal STR loci in a large population from Mainland China

**DOI:** 10.1101/015875

**Authors:** Zhuo Zhao, Hua Wang, Jie Zhang, Zhi-Peng Liu, Ming Liu, Yuan Zhang, Li Sun, Hui Zhang

## Abstract

STR, short trandem repeats, is well known as a type of powerful genetic marker and widely used in studying human population genetics. Compared with the conventional genetic markers, the mutation rate of STR is higher. Additionally, the mutations of STR loci do not lead to genetic inconsistencies between the genotypes of parents and children; therefore, the analysis of STR mutation is more suited to assess the population mutation. In this study, we focused on 15 autosomal STR loci (D8S1179, D21S11, D7S820, CSF1PO, D3S1358, TH01, D13S317, D16S539, D2S1338, D19S433, vWA, TPOX, D18S51, D5S818, FGA). DNA samples from a total of 42416 unrelated healthy individuals (19037 trios) from the population of Mainland China collected between Jan 2012 and May 2014 were successfully investigated. In our study, the allele frequencies, paternal mutation rates, maternal mutation rates and average mutation rates were detected in the 15 STR loci. Furthermore, we also investigated the relationship between paternal ages, maternal ages, pregnant time, area and average mutation rate. We found that paternal mutation rate is higher than maternal mutation rate and the paternal, maternal, and average mutation rates have a positive correlation with paternal ages, maternal ages and times respectively. Additionally, the average mutation rates of coastal areas are higher than that of inland areas. Overall, these results suggest that the 15 autosomal STR loci can provide highly informative polymorphic data for population genetic assessment in Mainland China, as well as confirm and extend the application of STR analysis in population genetics.

## Introduction

STRs are abundant in human genome and highly polymorphic due to allelic variations in the number of repeat units of 2-5 base pairs. For the reason of the susceptible slippage events during DNA replication, STR plays an important role as very useful genetic markers in the population genetics, linkage analysis, individual identification and parentage testing (Bar et al. 1997). As is well known, mutations of STR are relatively common in human genome. Compared with some other mutations in the genome, the mutation of STR has higher genetic diversity. Therefore, the mutation rate of STR has been commonly used in the assessment of population mutation associated with age, times, geographic regions and so on.

Previously, some reports have studied the mutation of STR in some areas and districts of China (Lei et al. 2014; Meng et al. 2015; Sun et al. 2014; Yan et al. 2006; Yin et al. 2015), but the scale of data on mutation rates of STR including the number of tested case, scope and area is still limited. Meanwhile, there are limited data of human population genetics from STR mutation. In this study, mutations of fifteen autosomal STR loci (D8S1179, D21S11, D7S820, CSF1PO, D3S1358, TH01, D13S317, D16S539, D2S1338, D19S433, vWA, TPOX, D18S51, D5S818, FGA) commonly used in parentage testing were studied. We first studied the frequency of all allele and mutation rate of STR locus. In addition, except for general analysis of average mutation rate, paternal mutation rate and maternal mutation rate, we further investigated the influence of repeat units of allele to the mutation of unit gains or losses. What’s more, we also investigated the relationship between paternal age, maternal age, pregnancy time, area and gamete’s mutation rate. Together, a total 19037 trios were checked for the biological consistency of paternity and maternity through analyses of fifteen autosomal STR loci. In our study, we finally observed total 678 mutations and to our best knowledge, this amounted STR mutation analysis is reported for the first time.

## Results

### Allele frequency and mutation model

We first investigated the frequencies of 15 autosomal STR loci and the results are shown in Figure 1 and supplementary Table 1. Generally, it is believed that the mutation of alleles occurs in the step wise mutation model (SMM) including single step mutation, double step mutation and multiple step mutations (Levinson and Gutman 1987). In our analysis, we observed one to four step mutations and a total number of 518 gains of repeat units compared with 439 losses of repeat units across all loci. Among these mutations, there are 487 1-step mutations, 135 2-step mutations, 58 3-step mutations, 18 4-step mutations and 53 unequivocal mutations in all loci. Mutations and the ratio of loss/gain repeat units of each allele of 15 STR loci were shown in Figure 2. We found that along with the increases of repeat unit, the ratio of loss/gain units increases gradually especially in THO1, D7S820 and D13S317 although the overall gains of repeats are higher than losses. Overall, Figure 2 not only presents a more detailed characterization of allele mutation of all loci, but also indentifies the relationship between the length of repeat units and the mutation of repeat losses or gains.

**Figure 1.**
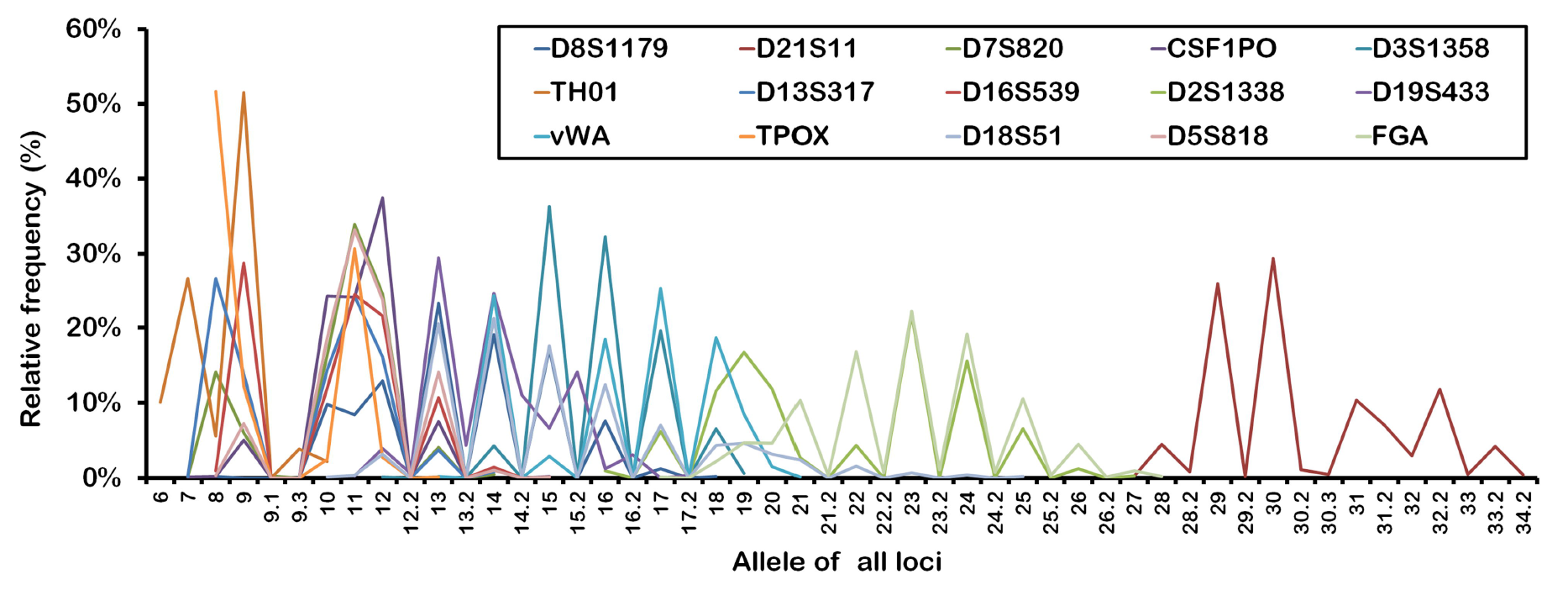
The distribution of allelic frequencies of 15 autosomal STR loci from china mainland population.

**Figure 2.**
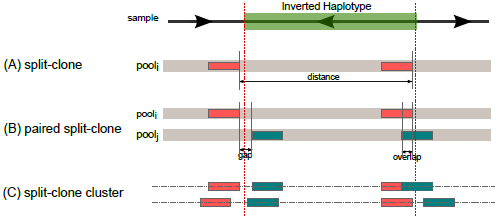
The mutation changes of repeats gains or losses of all allele in 15 STR loci. For each STR loci, the radio of lost number/gained number of repeat units was calculated. The change of color represents the changes of radio of loss/gain. Red: 100% of loss; Green: 100% of gain.

### Mutation rate of 15 autosomal STRs in mainland china

Next, we assessed the mutation rates including paternal mutation rate, maternal mutation rate and ambiguous. There are 289 mutations of paternity, 191 mutations of maternity and 198 ambiguous mutations (shown in Fig. 3 and supplementary Table 2). From the result, we found that the mutation rate of 15 STR loci was 0.05% to 0.63% and the highest mutation ratio was occurred in D13S317 loci and the lowest ratio in TPOX loci. Interestingly, across all loci, the paternal mutation rate was distinctly higher than the maternal mutation rate except for D19S433 (the paternal mutation rate and maternal mutation rate is roughly equal; paternity: 0.08%; maternity: 0.09%) (Fig. 3A) and the average ratio of male/female is approximately 1.43:1 (Fig. 3B, *: p<0.01). Sun *et al.* reported that the paternal mutations are more than maternal mutations in Guangdong Province of China and even the average ratio of male/female mutations can reach 3.8:1 (Sun et al. 2014). Our results are consistent with her conclusion.

**Figure 3.**
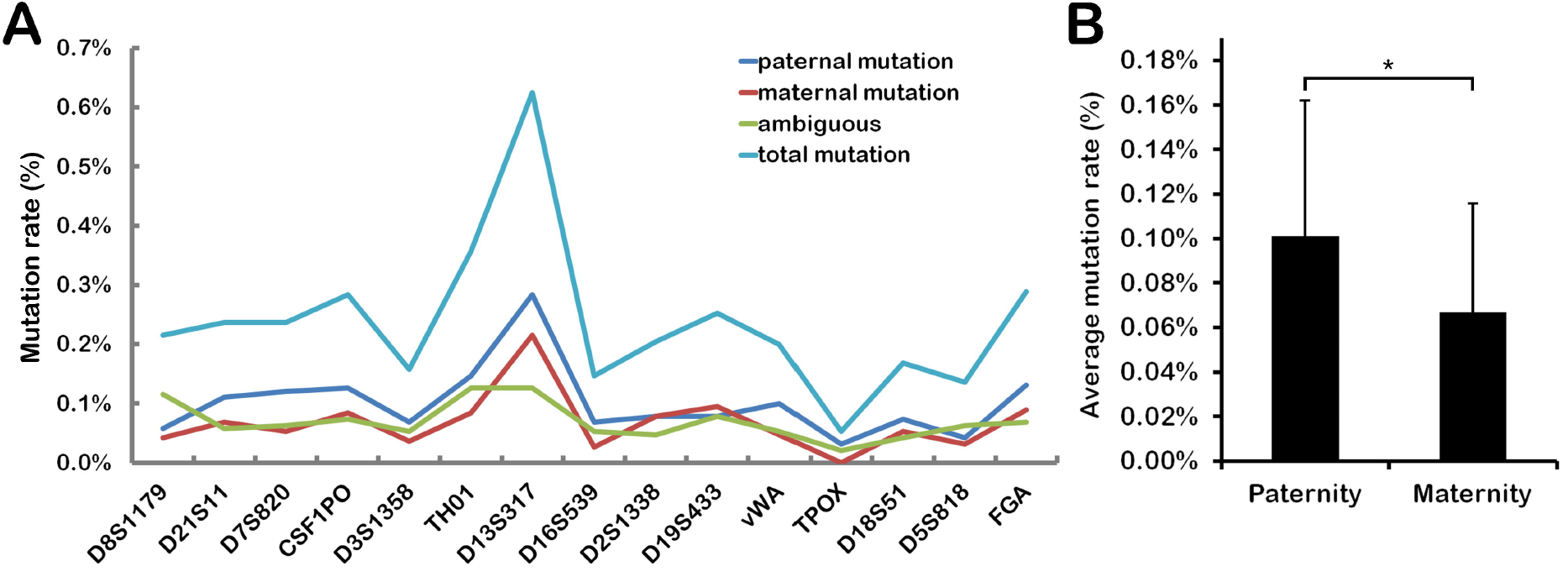
Mutated source analysis of children mutation in 15 STR loci. Observed mutations from 19037 trios were divided into paternal mutation, maternal mutation and ambiguous according to the mutation source. (A) The rate of paternal mutation, maternal mutation, ambiguous and total mutation at each STR loci. (B) The average mutation rate of all STR loci from paternity and maternity.

### Influence of parental age and pregnancy time on children’s mutation rates

In accordance with the information parental age and pregnancy time, we further studied the effects of them to children’s mutation and results were shown in Figure 4 and Figure 5. We found that there was a positive relationship between parental age and children’s mutation rate. With the increase of parental age, the children’s mutation rates have also increased (Fig. 4A, C) and especially with the increases of paternal age. The mutation rate of the research subjects with paternal age of ≥31years is higher than that of 25-30 years and ≤24 years (Fig. 4B). In our study, the influence of pregnancy time of maternity on children’s mutation was investigated too. The range of pregnancy time in our investigated case is from 1967 to 2013. Because there is a little amount of trios before 1987, according to the range of times trios were divided to four groups including group of before 1996, group of 1997-2001, group of 2002-2006 and group of after 2007. All loci mutation rate was analyzed and data was shown as Figure 5A. From the result, we found that times is earlier, the mutation rate of child is higher especially after 2002. Average mutation rate of all loci was calculated too and the result was shown as Figure 5B. Being consistent with Figure 5A, Figure 5B characterized a positive relationship between pregnancy time and mutation rate and moreover, after 2002 the mutation rate rises apparently.

**Figure 4.**
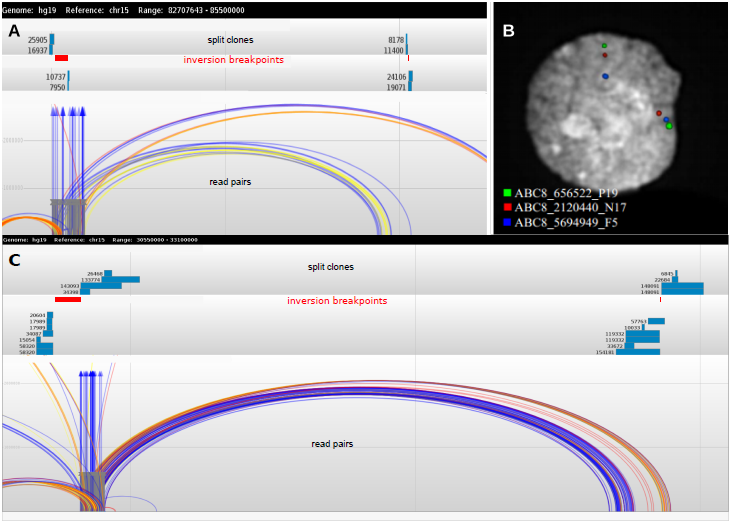
The relationship between parentage age and mutation rate. The average mutation rate of all loci at the certain age of paternity and maternity was analyzed. (A) The relationship between paternal mutation rate and paternal age. (C) The relationship between maternal mutation rate and maternal age. (B) All
paternal mutation events were divided into three groups according to paternal age including the group of ≤ 24, the group of 25-30 and the group of ≥31 and the average mutation rate of each groups was calculated.

**Figure 5.**
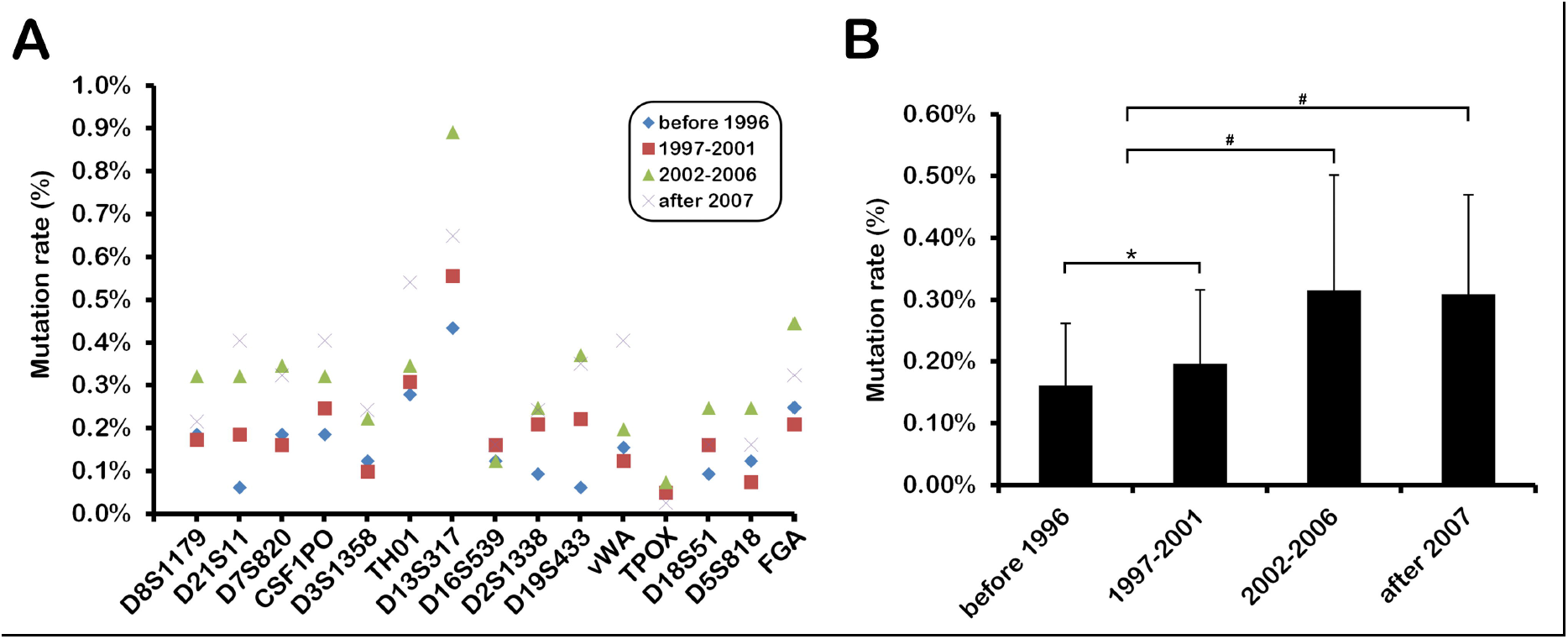
The relationship between pregnancy time and mutation rate. All mutation of 15 STR loci was divided into four groups according to the pregnancy time of maternity and for each loci, the average mutation rate of each group was calculated (A). (B) The average mutation rate of all loci was shown in the certain range of years.

### Area mutation rate in mainland china

The studied population located in Mainland china and distributed in most of province. In accordance with the mutation events in different province, we calculated the average mutation rate in mainland china area and data was shown as Figure 6. We divided the map of mainland china into six regions including Northeast China, East China, North China, South China, Southwest China and Northwest China and analyzed the mutation rates respectively in above region (Fig. 6A). We found that the mutation rates of Northeast China, East China, North China and South China were higher, but the mutation rates of Southwest China and Northwest China were lower. Mutation presents a tendency of that the population of east area has a higher mutation rate and the population of west area has a low mutation rate. The average mutation rate of east area, midland area and west area of mainland china was investigated too and result was consistent with Figure 6A (Fig. 6B).

**Figure 6.**
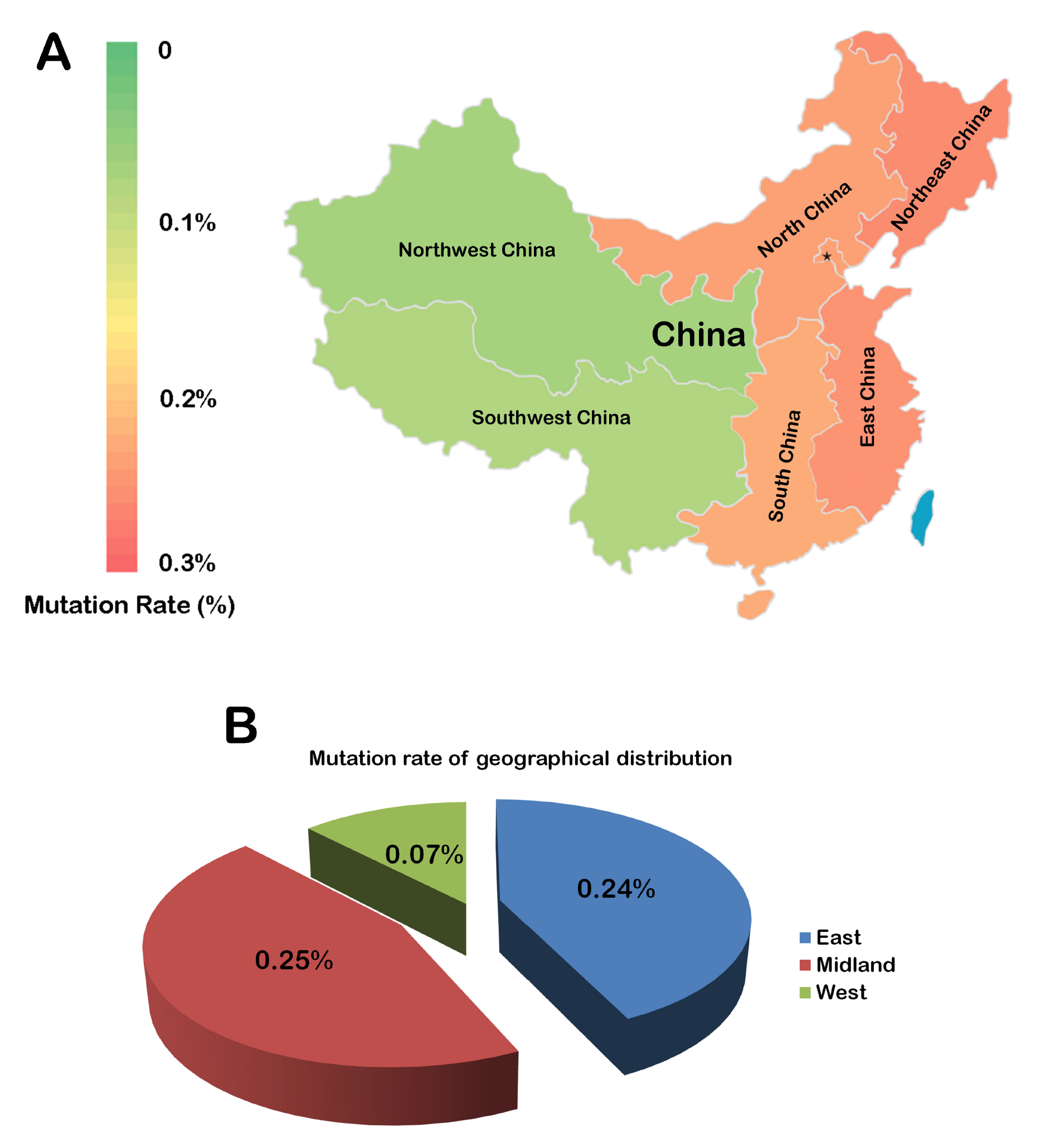
Distribution of mutations in the china mainland population. The area of china mainland was divided into six regions and included the regions of Northeast China, East China, North China, South China, Northwest China and Southwest China respectively. All mutations of above regions was investigated and the average mutation rate was shown (A). Red: high mutation rate; Green: low mutation rate. (B) The mutation of area of east, midland and west was respectively investigated and the mutation rate was shown.

## Discussion

Due to the high genetic polymorphism, STR has been used in human population studies as a genetic marker. In our study, we assessed 15 autosomal STR mutations from mainland china population through a large cohort (19037 trios).

First, all allele frequency and the mutation model from 15 STR loci were analyzed. We found that the mutation model of STR loci was mainly through repeat unit gains or losses. It is now generally accepted that replication slippage is the major mechanism causing STR allele mutation (Levinson and Gutman 1987). In our research, four types of allele mutation (1, 2, 3, 4-step mutation) including 698 mutated individuals were observed and these typed mutation accounted for about 93% of total mutations and our findings were accord with the stepwise mutation model. Besides, 53 unequivocal mutations were also observed and these mutation presented nonholonomic gain or loss of repeat units. These findings confirmed and expanded the mutation model of autosomal STR loci. What’s more, we found that there was a positive correlation between repeat losses and the number of repeat units. Along with the increases of repeat unit’s length, the ratio of lost units/gained units indicating that the repetitions of repeat unit could influence the mutation of losses. To our best knowledge, the effects of repetitions on loss mutation are first reported.

Taking the mutation source, paternal age and maternal age, all mutation events were analyzed. Our result showed that the paternal mutation is higher than maternal mutation and the parent age could increase the mutation rate on children’s STR loci. As we know, with the increases of mitosis the probability of mutation also increases (Fleming 1988; He et al. 2005; Riparbelli et al. 2004). In humans, the sperm mitosis is far more frequent than eggs mitosis, and therefore the paternal mutation should be higher in theory. In addition, it has been reported that estrogen plays an important role in the antioxidant activity against free radicals, mitochondria protection and anti-aging compared with androgen (Leal et al. 1998; Ruan et al. 2014; Stirone et al. 2005; Wang et al. 2014; Zetterberg and Celojevic 2015) indicating that estrogen might have a stronger protection. These finds can explain our results. In our study, we found that the average mutation rate from children of older parents is higher than younger parents, especially after 38 years (paternal age or maternal age). As is well known, telomerase play a crucial role in maintenance of telomere length and the stability of chromatin structure. Some reports showed that with age, the enzyme activity of telomerase will be decreased and telomerase gene might be mutated (Aubert and Lansdorp 2008; Dai et al. 2015; Hosen et al. 2014; Labussiere et al. 2014; Liu et al. 2014). These effects of telomerase can influence the length of telomere, induce mutation and cause tumorigenesis. Our findings about mutations mediated by parental age were consistent with these reports. Additionally, among the mutation influenced by paternal age and maternal age, the effects on mutation rate is stronger in paternal age than maternal age. We considered that the effective influence of paternity on children’s mutation might be due to that there is a difference of living habit, living environment and stress state between male and female.

The genetic polymorphism of some province of mainland china has been reported from many studies (Gao et al. 2014; He and Guo 2013; Hou et al. 2013; Sun et al. 2014; Wang et al. 2014; Wang et al. 2013; Weng et al. 2013; Zhang et al. 2014) and the mutation of population from some area of mainland has been also analyzed by some researcher (Brinkmann et al. 1998; Weng, Liu, Liu, Li, Liu, Wang 2013; Yan, Liu, Tang, Zhang, Huo, Hu, Yu 2006; Zhao et al. 2007). However, little is known about population genetics and mutation of whole mainland china. Our present work firstly describe the average mutation rate of 15 autosomal STR loci in Northwest China, East China, North China, South China, Southwest China, Northwest China area. Additionally, we have also investigated the relationship between the pregnancy time of maternal and average mutation rate. After performing the reform and opening-up policy, chinese economy has been a rapid growth and the living standard of people has been constantly improved. Nevertheless, with the rapid development of economy, the environmental pollution and the chance of toxic exposure have been more increased. From the point of view of area and time, the economy of east costal area is better than that of west inland area; moreover, china has entered into the phase of primary products production after 1995 and entered into the early industrialized after 2004 (Fu and Huang 2008; Liu et al. 2004; Qi et al. 2013; Xu and Li 2006). We considered that our result about the difference of mutation rate in area and times could be explain according to the development of chinese economy and the deterioration of living environment.

Taking together, our present studies are first to systematically characterize the population mutation through a large population in mainland china by using 15 auto STR mutation analysis. We hope that these STRs genetic polymorphism in china population could contribute to the analysis of its genetic diversity for the purpose of forensic testing or population genetic research.

## Material and Methods

Blood samples of unrelated healthy individual from 2012-2014 in the mainland china were collected. These individuals include 42, 416 DNA confirmed Chinese paternity testing case and in which, there are 19, 037 trios. In our study, the 19, 037 trios were as our research object. DNA from fresh blood was extracted and purified using PrepFiler™ Forensic DNA Extraction Kit (Applied Biosystems, Foster, CA, USA) and amplification of the STRs was performed using the AmpFlSTR® Identifiler® PCR Amplification Kit (Applied Biosystems, Foster, CA, USA) according to the manufacture instructions. The kits include 15 STR loci (D8S1179, D21S11, D7S820, CSF1PO, D3S1358, TH01, D13S317, D16S539, D2S1338, D19S433, vWA, TPOX, D18S51, D5S818, FGA) and a sex determination gene (Amelogenin). STR genotyping was done on the ABI PRISM^®^ 3130xl Genetic Analyzer (Applied Biosystems, Foster City, CA, USA) and analyzed using the GeneMapper ID 3.2 software (Applied Biosystems, Foster City, CA, USA). All experiments were carried out according to the kit control and laboratory internal control standards.

Null alleles were assumed and counted in cases of single discrepancies between a homozygous parent and a homozygous child at a locus when at least 12 other independent autosomal STR loci were consistent with paternity and/or maternity. Mutations were assumed and counted after excluding the null alleles in cases of single discrepancies between a parent and child at a locus when at least 12 other STR loci were consistent with paternity and/or maternity.

The experimental data was shown as the mean ± standard deviation. Statistical significance was evaluated with SPSS software. Experimental differences were analyzed using the two-tailed Student’s t-text. P values<0.05 was considered statistic significant.

## Acknowledgment

We are grateful to TIANJIN JINSHI FORENSIC CENTER for the support of informations.

## References

1. Aubert G, Lansdorp PM. 2008. Telomeres and aging. Physiol Rev 88: 557–579.

2. Bar W, Brinkmann B, Budowle B, Carracedo A, Gill P, Lincoln P, Mayr W, Olaisen B. 1997. DNA recommendations. Further report of the DNA Commission of the ISFH regarding the use of short tandem repeat systems. International Society for Forensic Haemogenetics. Int J Legal Med 110: 175–176.

3. Brinkmann B, Klintschar M, Neuhuber F, Huhne J, Rolf B. 1998. Mutation rate in human microsatellites: influence of the structure and length of the tandem repeat. Am J Hum Genet 62: 1408–1415.

4. Dai J, Cai H, Zhuang Y, Wu Y, Min H, Li J, Shi Y, Gao Q, Yi L. 2015. Telomerase gene mutations and telomere length shortening in patients with idiopathic pulmonary fibrosis in a Chinese population. Respirology 20: 122–128.

5. Fleming AF. 1988. Possible aetiological factors in leukaemias in Africa. Leuk Res 12: 33–43.

6. Fu M, Huang ZM. 2008. The Kuznets relationship between economic development stages and environmental pollution in China. China Industrial Economics 35–43.

7. Gao Y, Han JT, Shen CM, Wu H, Yuan GL, Zhao LJ, Yan JW, Meng HT, Zhang YD, Liu WJ, et al. 2014. Structural polymorphism analysis of Chinese Mongolian ethnic group revealed by a new STR panel: genetic relationship to other groups. Electrophoresis 35: 2008–2013.

8. He J, Guo F. 2013. Population genetics of 17 Y-STR loci in Chinese Manchu population from Liaoning Province, Northeast China. Forensic Sci Int Genet 7: e84–e85.

9. He N, Li C, Zhang X, Sheng T, Chi S, Chen K, Wang Q, Vertrees R, Logrono R, Xie J. 2005. Regulation of lung cancer cell growth and invasiveness by beta-TRCP. Mol Carcinog 42: 18–28.

10. Hosen I, Rachakonda PS, Heidenreich B, Sitaram RT, Ljungberg B, Roos G, Hemminki K, Kumar R. 2014. TERT promoter mutations in clear cell renal cell carcinoma. Int J Cancer [Epub ahead of print].

11. Hou G, Jiang X, Wang Y, Li Q, Sun H. 2013. Genetic distribution on 15 STR loci from a Han population of Shenyang region in northeast China. Forensic Sci Int Genet 7: e86–e87.

12. Labussiere M, Di Stefano AL, Gleize V, Boisselier B, Giry M, Mangesius S, Bruno A, Paterra R, Marie Y, Rahimian A, et al. 2014. TERT promoter mutations in gliomas, genetic associations and clinico-pathological correlations. Br J Cancer 111: 2024–2032.

13. Leal AM, Begona RM, Martinez R, Lacort M. 1998. Cytoprotective actions of estrogens against tert-butyl hydroperoxide-induced toxicity in hepatocytes. Biochem Pharmacol 56: 1463–1469.

14. Lei L, Xu J, Du Q, Fu L, Zhang X, Yu F, Ma C, Cong B, Li S. 2014. Genetic polymorphism of the 26 short tandem repeat loci in the Chinese Hebei Han population using two commercial forensic kits. Mol Biol Rep 42: 217–225.

15. Levinson G, Gutman GA. 1987. Slipped-strand mispairing: a major mechanism for DNA sequence evolution. Mol Biol Evol 4: 203–221.

16. Liu T, Wang N, Cao J, Sofiadis A, Dinets A, Zedenius J, Larsson C, Xu D. 2014. The age- and shorter telomere-dependent TERT promoter mutation in follicular thyroid cell-derived carcinomas. Oncogene 33: 4978–4984.

17. Liu XH, Wang JF, Meng B. 2004. Analysis on China’s spatio-temporal dynamics and imbalance of regional economy. Geographical Research 23: 530.

18. Meng HT, Zhang LP, Wu H, Yang CH, Chen JG, Wang Y, Yan JW, Wang HD, Zhang YD, Liu WJ, et al. 2015. Genetic diversities of 20 novel autosomal STRs in Chinese Xibe ethnic group and its genetic relationships with neighboring populations. Gene 557: 222–228.

19. Qi YJ, Yang Y, Jin FJ. 2013. China’s economic development stage and its space-time evolution characteristics. ACTA GEOGRAPHICA SINICA 68: 517–531.

20. Riparbelli MG, Massarelli C, Robbins LG, Callaini G. 2004. The abnormal spindle protein is required for germ cell mitosis and oocyte differentiation during Drosophila oogenesis. Exp Cell Res 298: 96–106.

21. Ruan Y, Wu S, Zhang L, Chen G, Lai W. 2014. Retarding the senescence of human vascular endothelial cells induced by hydrogen peroxide: effects of 17beta-estradiol (E2) mediated mitochondria protection. Biogerontology 15: 367–375.

22. Stirone C, Duckles SP, Krause DN, Procaccio V. 2005. Estrogen increases mitochondrial efficiency and reduces oxidative stress in cerebral blood vessels. Mol Pharmacol 68: 959–965.

23. Sun H, Liu S, Zhang Y, Whittle MR. 2014. Comparison of southern Chinese Han and Brazilian Caucasian mutation rates at autosomal short tandem repeat loci used in human forensic genetics. Int J Legal Med 128: 1–9.

24. Sun M, Zhang X, Wu D, Shen Q, Wu Y. 2014. Population genetic data of 15 STR loci in Gansu Han population from China. Int J Legal Med 9: 30–32.

25. Wang F, Xiao J, Shen Y, Yao F, Chen Y. 2014. Estrogen protects cardiomyocytes against lipopolysaccharide by inhibiting autophagy. Mol Med Rep 10: 1509–1512.

26. Wang HD, Ruan JG, Shen CM, Zhang YD, Liu WJ, Meng HT, Yang XY, Yan JW, Liao SX, Fan SL, et al. 2014. Polymorphic distribution and forensic effectiveness study of eight miniSTR in Chinese Uyghur ethnic group. Mol Biol Rep 41: 2371–2375.

27. Wang XQ, Wang CC, Deng QY, Li H. 2013. [Genetic analysis of Y chromosome and mitochondrial DNA poly-morphism of Mulam ethnic group in Guangxi, China]. Yi Chuan 35: 168–174.

28. Weng W, Liu H, Liu C, Li S, Liu C, Wang H. 2013. [Haplotype frequency and mutation of 17 Y-STR loci in Han population in Guangdong Province]. Nan Fang Yi Ke Da Xue Xue Bao 33: 412–415.

29. Xu ZY, Li ST. 2006. Analysis on the trend of regional income disparity in China. Economic Research Journal: 106–116.

30. Yan J, Liu Y, Tang H, Zhang Q, Huo Z, Hu S, Yu J. 2006. Mutations at 17 STR loci in Chinese population. Forensic Sci Int 162: 53–54.

31. Yin C, Ji Q, Li K, Mu H, Zhu B, Yan J, Yu Y, Wang J, Chen F. 2015. Analysis of 19 STR loci reveals genetic characteristic of eastern Chinese Han population. Forensic Sci Int Genet 14: 108–109.

32. Zetterberg M, Celojevic D. 2015. Gender and cataract - the role of estrogen. Curr Eye Res 40: 176–190.

33. Zhang L, Zhao Y, Guo F, Liu Y, Wang B. 2014. Population data for 15 autosomal STR loci in the Miao ethnic minority from Guizhou Province, Southwest China. Forensic Sci Int Genet.

34. Zhao ZM, Liu Y, Lin Y. 2007. [Mutations of short tandem repeat loci in Identifiler system]. Fa Yi Xue Za Zhi 23: 290–291, 294.

